# Sample preparation and warping accuracy for correlative multimodal imaging in the mouse olfactory bulb using 2-photon, synchrotron X-ray and volume electron microscopy

**DOI:** 10.1101/2022.02.18.481045

**Authors:** Yuxin Zhang, Tobias Ackels, Alexandra Pacureanu, Marie-Christine Zdora, Anne Bonnin, Andreas T. Schaefer, Carles Bosch

## Abstract

Integrating physiology with structural insights of the same neuronal circuit provides a unique approach to understanding how the mammalian brain computes information. However, combining the techniques that provide both streams of data represents an experimental challenge. When studying glomerular column circuits in the mouse olfactory bulb, this approach involves e.g. recording the neuronal activity with *in vivo* 2-photon (2P) calcium imaging, retrieving the circuit structure with synchrotron X-ray computed tomography with propagation-based phase contrast (SXRT) and/or serial block-face electron microscopy (SBEM) and correlating these datasets. Sample preparation and dataset correlation are two key bottlenecks in this correlative workflow. Here, we first quantify the occurrence of different artefacts when staining tissue slices with heavy metals to generate X-ray or electron contrast. We report improvements in the staining procedure, ultimately achieving perfect staining in ∼67% of the 0.6 mm thick olfactory bulb slices that were previously imaged *in vivo* with 2P. Secondly, we characterise the accuracy of the spatial correlation between functional and structural datasets. We demonstrate that direct, single-cell precise correlation between *in vivo* 2P and SXRT tissue volumes is possible and as reliable as correlating between 2P and SBEM. Altogether, these results pave the way for experiments that require retrieving physiology, circuit structure and synaptic signatures in targeted regions. These correlative function-structure studies will bring a more complete understanding of mammalian olfactory processing across length scales and time.

## Introduction

Maps are essential tools to accelerate discovery across charted and uncharted domains. By providing information in an organised and intuitive structure, they can guide the development of strategic approaches that maximise gain while minimising risk. As it has occurred in a myriad of disciplines before, maps are also playing a crucial role in supporting research in the life sciences and in neuroscience in particular (Niedworok et al., 2016; Winnubst et al., 2019; Claudi et al., 2020; Ueda et al., 2020; BRAIN Initiative Cell Census Network (BICCN), 2021). Mapping the brain at multiple scales can help gain mechanistic insights into its function that might already be present in existing data, as well as guide the exploration of the uncharted territories that hold the keys for unsolved functions. Creating those maps requires recording the spatial distribution of relevant brain properties across a volume that encodes functional units at a resolution that can discriminate between individual operators. In the mammalian brain, neuronal circuits usually span above 1 mm^3^, while the dimensions of functionally relevant operators (nuclei, dendrites, axons, synapses) range between 10 µm - 10 nm (Helmstaedter, 2013; Klinger et al., 2021). Targeted maps carrying more detail (only) where it is needed offer a valuable solution to this vast challenge, and can be generated by means of correlative multimodal imaging (CMI) pipelines (Bock et al., 2011; Briggman et al., 2011; Karreman et al., 2016; Lee et al., 2016; Dyer et al., 2017; Bosch et al., 2021). The reliability of those brain maps will be directly influenced by two main factors: the quality of the images of each individual modality and the quality of their registration into a multimodal map.

The quality of an image is limited by multiple factors including optics, detector sensitivity, exposure time or the structure of the specimen leading to signal contrast. The latter is engineered and optimised by preparing the specimen for a particular imaging technique in a process often referred to as the sample preparation protocol (SPP). An SPP incorporates all transformations necessary to make a specimen compatible with an imaging technique and may involve mechanical procedures (such as slicing), chemical transformations (such as fixing) and signal transformations (such as staining). SPPs can be studied as a sequence of smaller-scale operations sharing a single goal, which we will refer to as ‘steps’. In that way, a single step may involve slicing a fresh piece of brain tissue to a specific thickness under controlled constraints of temperature, osmolarity and time, while another might involve chemically staining a tissue slice with a buffered solution of osmium tetroxide while controlling for temperature, duration and agitation. Optimising an SPP presents therefore a multivariate challenge that can be approached by making informed decisions to optimise each step individually while monitoring global yields.

The maps guiding our understanding of how neuronal circuits operate in the mammalian brain should convey spatial and functional information across the critical dimensions specified above and can be obtained using a CMI pipeline that combines *in vivo* 2-photon (2P) microscopy of intracellular [Ca^2+^], synchrotron X-ray computed tomography with propagation-based phase contrast (SXRT) and serial block-face electron microscopy (SBEM) (Bosch et al., 2021). The SPP for this approach will therefore need to be compatible with *all* techniques involved and its optimal implementation will be the one that accomplishes two main requisites: (1) it delivers an optimal final map, therefore balancing inter-modality trade-offs towards the benefit of the initial research question; and (2) it provides the highest yield, making it compatible with more challenging research questions in the future.

The datasets carrying the best spatial resolution in this CMI pipeline will be provided by SBEM (Bosch et al., 2021). In these, the signal is generated by recording the back-scattered electrons arising from a 1-2 keV electron beam scanning the surface of a polished tissue block at an electron dose of typically <20 e^-^/nm^2^ (Kubota, 2015). For soft tissue to generate contrast under this regime, tissue is stained with heavy metals (Hayat, 2000). Heavy metals bind non-specifically to lipids and proteins, ultimately providing a sample with a topology that faithfully reproduces its ultrastructure (Buckley and Porter, 1967). SBEM adds one challenge to otherwise more conventional transmission electron microscopy imaging: here, part of the electron beam will be absorbed by the sample, locally increasing its charge and potentially influencing the trajectories of neighbouring back-scattered electrons. In order to prevent such charging artefacts, SBEM-specific SPP optimisations have incorporated elements that increase the conductivity of the sample -such as by increasing the total metal load employed in staining, in terms of both the number of metallic compounds and their concentrations (Tapia et al., 2012). The diffusion mechanism used to deliver those metals through the tissue remains passive, and therefore incubation time is subject to a quadratic relationship with the distance each compound should travel (Ströh et al., 2021), imposing severe temporal burdens for staining specimens of dimensions beyond 1 mm (Mikula and Denk, 2015). Moreover, abnormalities arising from individual staining steps can interact with each other (Hua et al., 2015; Mikula and Denk, 2015), creating an additional challenge to ensure all compounds optimally stain the tissue across all dimensions. Nevertheless, some SPPs have reported acceptable SBEM results in mouse brain tissue samples of thicknesses ranging from 0.1 mm to a complete mouse brain (∼10 mm) (Briggman et al., 2011; Tapia et al., 2012; Wilke et al., 2013; Hua et al., 2015; Mikula and Denk, 2015; Pallotto et al., 2015; Wanner et al., 2016; Genoud et al., 2018), providing a promising starting point for further optimisations aiming to obtain a multimodal brain map. Interestingly, such SBEM staining also provides good contrast in SXRT (Bushong et al., 2015) because X-ray absorption is proportional to atomic number. Hence heavy metal-stained samples of thicknesses <1 mm can be imaged efficiently with this technique.

Finally, the registration quality of the individual map layers needs to be accurate enough to correlate any feature of interest across all modalities. For maps aimed at pinpointing the structure of a neural circuit, registration should be accurate at single-cell precision. Since the largest feature to unequivocally identify a single cell within a circuit is its nucleus, the requirement becomes registering at a precision better than the inter-nucleus distance, which is approximately 20 µm in the mitral cell layer of the olfactory bulb (Panhuber et al., 1985; Frazier and Brunjes, 1988; Parrish-Aungst et al., 2007).

Here we show an optimization of a CMI pipeline aiming to map a mammalian brain circuit with 2P microscopy, SXRT and SBEM. Using the mouse olfactory bulb, we assessed and made improvements on the SPPs and quantified the registration accuracy across imaging modalities. We demonstrate that this pipeline is effective and that single-cell registration accuracy can be readily achieved between 2P and SXRT.

## Materials and methods

All SPPs used in this study are described in detail in **Supp. Table 2** and are available for further analysis (see **Data availability statement**). The experimental details of samples shown in the figures are listed in **Supp. Table 3**.

### Animals

All animals used were 8-24 week-old mice of C57/Bl background and mixed gender. Details of the transgenic mouse lines are listed in **Supp. Table 4**. All animal protocols were approved by the Ethics Committee of the board of the Francis Crick Institute and the United Kingdom Home Office under the Animals (Scientific Procedures) Act 1986.

### Craniotomy and durectomy surgery

In experiments involving *in vivo* 2-photon (2P) imaging, anaesthesia was induced and maintained with intraperitoneal injection of sleep mix (0.05 mg/kg fentanyl, 5 mg/kg midazolam, 0.5 mg/kg medetomidine) and its depth monitored by toe-pinch reflex tests. Body temperature of mice was measured with a rectal temperature probe and maintained at 36-38 °C with a thermoregulator (DC Temperature Controller, FHC, ME USA) heat pad.

Following shaving, skin disinfection with 1% chlorhexidine and skin incision over the skull, a custom head-fix implant with a central hole was fixed to the skull with medical super glue (Vetbond, 3M, Maplewood MN, USA) and dental cement (Paladur, Heraeus Kulzer GmbH, Hanau, Germany; Simplex Rapid Liquid, 437 Associated Dental Products Ltd., Swindon, UK) so that the nasal bone above olfactory bulbs were exposed through the headplate hole. A ∼2 mm-diameter craniotomy over the left olfactory bulb was made with a dental drill (Success 40, Osada, Tokyo, Japan). ACSF (10 mM HEPES, 10 mM glucose, 125 mM NaCl, 5 mM KCl, 2 mM MgSO_4_.7H_2_O, 2 mM CaCl_2_.2H_2_O, pH = 7.4) was used to moisten the brain throughout the surgery. Exposed dura was removed with forceps and needles. Then a drop of warm 2% low-melt agarose diluted in ACSF was applied, quickly followed by the placement of a glass window cut from a coverslip (borosilicate glass #1 thickness (150 μm)). The edge of the window was glued to the nasal bones with medical super glue (Vetbond, 3M, Maplewood MN, USA).

Some SPPs also include measures to prevent the formation of edema, including the use of ice-cold ACSF during surgery to cool down the skull from drilling and the use of dexamethasone (20 μl/mouse intraperitoneal injection) 2-4h prior to surgery or dissection (**Supp. Table 2**).

### In vivo 2-photon (2P) imaging

Following surgery, animals were transferred to a 2P microscope (Scientifica Multiphoton VivoScope coupled with a MaiTai DeepSee Laser tuned to 910-940 nm, <30 mW underneath objective) and held by a custom head-fix implant holder. Respiration was measured at the right nostril with a flow sensor (A3100, Honeywell, NC, USA) and the signal was digitised with a Power 1401 ADC board (CED, Cambridge, UK). Odours were delivered at inhalation onset through custom-made olfactometers (Ackels et al., 2021; Bosch et al., 2021) whose output port was <1 cm away from the left nostril. Images (512×512 pixels, field of view 550×550 μm) were taken with a resonant-galvo scanner at 30 Hz frame rate. Six or twelve planes covering ∼300 μm in z, including the mitral cell layer, were acquired with a 16x 0.8 NA water-immersion objective (Nikon) coupled to a piezo motor (PI Instruments, UK). The fluorescently labelled glomerulus was used to position the field-of-view.

Following functional imaging, blood vessels were fluorescently labelled by injecting the tracer sulforhodamine 101 intraperitoneally (50 µl of a 0.1 M solution) (Sigma Aldrich). A dataset with voxels sized 0.358 µm in x, y and 5 µm in z containing the entire functionally-recorded volume was then acquired. Red and green channels were used to capture the vasculature and the resting GCaMP signals, respectively.

### Dissection, slicing and fixation

For immersion fixation, animals were culled by cervical dislocation or decapitation after intraperitoneal injection of an overdose of pentobarbital (600 mg/kg pentobarbitone with mepivacaine) or sleep mix (0.05 mg/kg fentanyl, 5 mg/kg midazolam, 0.5 mg/kg medetomidine). Ice-cold dissection solution (4.6% sucrose, 0.563 mM CaCl_2_, 65.2 mM NaH_2_PO_4_.H_2_O, with 0.02% sodium azide, bubbled with 95% O_2_/5% CO_2_, osmolarity 280-320 mOsm/l) was used both to keep tissue cold during dissection and as the bathing solution during vibratome slicing (Leica VT1200S). 0.4-1 mm-thick dorsal OB slices were taken by near-horizontal slicing, setting the vibratome blade tangent to the OB surface. Half and whole OB samples were cut with a scalpel. Specimens were then transferred to cold fixative (see below) and post-fixed overnight at 4°C.

For perfusion, animals were anaesthetized with an overdose of pentobarbital (600 mg/kg pentobarbitone with mepivacaine) administered intraperitoneally. The skin over the chest was removed and incisions made medially-laterally across the abdomen muscle and across the diaphragm. Before cardiac arrest took place, room temperature fixative (see below) was delivered by a cannula inserted into the left atrium of the heart driven by a peristaltic pump (4.5-7.2 ml/min, 80-100 ml). The brain was dissected out and post-fixed overnight at 4°C and OBs vibratome-sliced on the following day in 0.15 M NCB (sodium cacodylate (aq), pH = 7.4, 280-320 mOsm/l).

The fixatives used were:

a. 1% or 2% glutaraldehyde (GA) with 2 mM CaCl_2_ and 0.02% sodium azide in 0.15 M sodium cacodylate buffer (NCB), based on (Pallotto et al., 2015).
b. 1.25% GA and 2.5% paraformaldehyde (PFA) or 2% GA and 2.5% PFA with 2 mM CaCl_2_ in 0.08 M NCB, based on (Hua et al., 2015).
c. 0.5% GA or 1.25% GA or 2% GA or 1% PFA or 2.5% PFA or 4% PFA with 2 mM CaCl_2_ in 0.08 M NCB.

### Epifluorescence and ex vivo 2P imaging

For samples containing a fluorescently-labelled glomerulus, after a brief contact with fixative, epifluorescence images tiling the entire specimen were recorded with a widefield fluorescence microscope (Olympus Axioplan2 with LEJ ‘ebq 100 isolated-z’ lamp) before overnight postfixation.

After postfixation, two volumes covering the fluorescently-labelled glomerulus (located using the epifluorescence images as guides) and its surrounding regions were acquired and stitched together (voxels sized 1.074 µm in x, y and 5 µm in z) with the aforementioned 2P system (laser wavelength 800 nm). This dataset featured glomeruli and surface blood vessels from the autofluorescence signal and was used to aid the relocation of functionally imaged volume in later acquired SXRT/EM datasets.

### Staining, dehydration and embedding

Fixed OB slices were washed in 0.15 M NCB (sodium cacodylate (aq.) with or without 0.02% sodium azide, pH = 7.4, 280-320 mOsm/l, 3×1h), then stained using an automated tissue processor (Leica EMTP). The full descriptions can be found in **Supp. Table 2**. In brief, specimens were processed with one of the following staining protocols:

a. 2% OsO_4_, 3% potassium ferrocyanide (KFe), 2 mM CaCl_2_ in 0.15 M NCB (1h, 4°C); 1% thiocarbohydrazide (TCH, aq., 1h, 50°C); 2% OsO_4_ (aq., 1h, 20°C); 2% uranyl acetate (UA, aq., overnight, 4°C); lead aspartate (aq. pH = 5.5, 2h, 60°C) with water washes between each step, protocol referred to as ‘**1sRO half time**’.
b. same as a) but doubled incubation times of all steps except UA, protocol referred to as ‘**1sRO**’.
c. 2% OsO_4_ in 0.15 M NCB (1.5h, 20°C) followed by 2.5% KFe in 0.15 M NCB (1.5h, 20°C) without water wash; 1% TCH (aq., 45min, 40°C); 2% OsO_4_ (aq., 1.5h, 20°C); 1% UA (aq.) warmed to 50°C for 2h after overnight at 4°C; lead aspartate (aq., pH = 5.0, 2h, 50°C) with water washes between each step, protocol referred to as ‘**2sRO**’.
d. same as c) but TCH (aq.) incubated for 1h at 50°C and lead aspartate (pH = 5.5) at 60°C, used in exp. A and exp. B in **Fig. 3b1**
e. same as c) but all warm incubations happening at 60°C and all steps performed manually. There were 6 groups of samples in this SPP, corresponding to terminating the staining at each of the 6 reagents (**Fig. 3d1**). Early finishing samples were pooled in water at room temperature while others finished their course of staining.

Samples were then dehydrated with 75%, 90%, 100%, 100% ethanol series, transferred to propylene oxide and infiltrated with 25%, 50%, 75%, 100%, 100% hard epon diluted in propylene oxide, and finally polymerized for 72h at 60-70°C. For experiments in **Fig. 3d1-2**, samples were dehydrated with 100% ethanol at 4°C overnight before being transferred to propylene oxide.

### LXRT imaging

After staining and embedding, the samples were imaged with a laboratory-source X-ray microscope (LXRT; Zeiss Xradia 510 Versa, 40 kV, 3W, LE2-4 filter, 1601 projections over 180° of sample rotation, 3-20s exposure time per frame, 1 tile, 3-6 μm pixel size). The reconstructed tomogram was used to screen for artefacts.

### SXRT imaging

The SXRT dataset analysed in this work is C525, described previously in (Bosch et al., 2021). In brief, samples were imaged with synchrotron X-ray computed tomography exploiting propagation-based phase contrast at the I13-2 beamline of the Diamond Lightsource (Didcot, UK; 22 keV, 3001 projections over 180° of sample rotation, 0.4 s exposure time per frame, 2-24 tiles, 325 nm isotropic voxel size). The reconstructed tomogram is referred to as the SXRT dataset.

### Trimming and SBEM imaging

To meet the sample size and conductance requirements of our electron microscopes, the resin embedded samples were further processed prior to electron microscopy. Firstly, trims were planned from rendered LXRT or SXRT datasets in Amira or Imaris. 2P datasets were warped into LXRT or SXRT space and overlaid on top to help target and preserve the functionally imaged volume in the trimmed block. With the help of this trimming plan, embedded samples were trimmed to a suitable shape and size with a razor blade and a glass and/or diamond knife (Diatome) at an ultramicrotome (Leica UC7). Subsequently, the sides of the sample were painted with silver colloidal suspension (SPI supplies) and sputter-coated with a layer of 10 nm-thick platinum (Quorum Q150R S).

2D images used for ultrastructure checks (**Supp. F3b**) were obtained in a 3View2 – Zeiss Merlin SBEM under high vacuum (1.5 kV, 0.2 nA, 2 us/pixel, 10 nm or 50 nm pixel size) using an OnPoint (Gatan) back-scattered electron detector.

The volume SBEM dataset analysed in this study comes from sample C525a, described previously in Bosch et al. (2021). This volume SBEM dataset was obtained with a 3View2 - variable pressure Zeiss Sigma serial-block face SEM using focal charge compensation (2.5 kV, 30 μm aperture, (80 nm)^3^ voxel size).

### Artefact categorization

After looking through all LXRT datasets, 12 types of artefacts were defined according to their appearance in the LXRT datasets (**Supp. Table 1**). For each sample, a score of 0 (absent) or 1 (present) was given to each artefact type and the following variables compiled: ‘experiment ID’, ‘experiment start date’, ‘fixation method’, ‘fixative formula’, ‘staining method’, ‘anaesthetics’, ‘culling method’, ‘use of dexamethasone’, ‘use of cold buffer’, ‘mouse line’, ‘tissue type’, ‘slice index’, ‘thickness’ and ‘2P history’. For overall occurrence rate quantification, all samples from all experiments were included. Other analyses were done using samples from relevant experiments (**Supp. Table 5**). Analyses were performed with Python3 (Van Rossum and Drake, 2009) in a Jupyter notebook (Kluyver et al., 2016). The artefact scoring tables and analysis scripts are available (see **Data availability statement**).

### Sample volume estimation

In total, the final ‘in resin’ volume of 80 samples was measured from their LXRT datasets (**Supp. Table 6**). Sample volume was measured as the number of voxels containing tissue times the voxel size. The voxels containing tissue data were masked in Fiji (Schindelin et al., 2012) by thresholding the signal and discarding low-signal background voxels. For samples with ‘crack’ artefact, the volume occupied by ‘crack’ was excluded from the tissue voxel count.

To estimate the volume of the fresh tissue samples, we utilised landmark pairs from the set of ‘warping landmarks’ used to calculate the warp field (see results and below). Paired landmarks mark the same features of a sample in both datasets. Changes in tissue volume after sample preparation will therefore be reflected in changes in distances between landmarks. For one sample, 20 landmarks from the 2P dataset were randomly selected and their distances to every other landmark calculated. The top 400 distances were taken (Df1_400). Distances between these landmarks were also calculated for the structural dataset of the same sample (Ds1_400). The amount of sample volume change (r) was taken to be the averaged ratio between Ds1_400 and Df1_400 cubed. The fresh sample volume is therefore the CT volume divided by r (**Supp. F4**).

### Tracing, warping and warping accuracy estimation

Dataset annotations, including somata seeds and blood vessel tracings, were generated with the data storage, display and annotation tool webKnossos (Boergens et al., 2017).

Warping landmarks were seeded manually using the Fiji plugin BigWarp (Bogovic et al., 2016) and then exported. Landmarks were any distinguishable features shared by the datasets involved, such as blood vessel branching points. The warping function was then calculated using the exported landmark file and a function from the webKnossos skeleton analysis toolbox built in the Matlab (Mathworks) environment. This toolbox is available online (see **Data availability statement**).

For the C525a dataset, the warping function between 2P and SBEM was calculated using 171 landmarks seeded throughout the dataset (**Fig. 6a**). Similarly, 64 and 51 landmarks were used for 2P to SXRT warping and SXRT to SBEM warping, respectively. These sets of paired landmarks were referred to as ‘warping landmarks’ in the results section.

For warping accuracy quantification, blood vessels in the upper EPL within a (160 μm)^3^ bounding box were traced in the 2P, SXRT and SBEM datasets of sample C525a. To trace the same sets of blood vessels in all three datasets, the bounding boxes used were centred on a functionally imaged soma in 2P and its warped location in SXRT and SBEM. Due to the resolution limit, thin blood vessels were not traceable in the SXRT dataset and the corresponding blood vessel tracings in the 2P and SBEM datasets were deleted for consistency. After this, all blood vessels branching points (junctions) were named and matched across the three datasets using blood vessel shape as a guide. Lastly, freely ending blood vessel tracings were trimmed up until a junction. Therefore, the same blood vessels were covered to the same extent in all three datasets and hence could be compared later. Processed tracings containing those annotations in the different datasets are available online (see **Data availability statement**).

The blood vessel tracings were then warped into a common framework (e.g. SBEM space, **Fig. 6c**). Each tracing was then split into segments at blood vessel junctions. For each segment, the average segment length (among 2P, SBEM and SXRT) was calculated. This length was then divided by an interpolation step size of 500 nm, giving the number of steps (n) to interpolate for this segment. Subsequently, each segment from each dataset was interpolated with n steps, giving n points on the blood vessel that are matched across all three datasets. The tracing accuracy at that location between any two datasets was then estimated to be the distance between the matched points (**Fig. 6d**). The overall warping accuracy between any two datasets was taken to be the mean of all matched-point distances.

## Results

The olfactory bulb contains the first neuronal circuit in the olfactory sensory pathway, the glomerular column (Mori et al., 1999). Thick (>0.4 mm) dorsal slices of the bulb contain all the histological layers embedding that circuit (**Fig. 1a**). We prepared 415 0.4-1mm-thick vibratome-cut slices of mouse olfactory bulb for electron microscopy (**Fig. 1a1**), of which 328 were first dorsal slices (**Fig. 1a2**). However, first slices are asymmetrical, with one side containing biological features potentially capable of acting as a barrier for the diffusion of the staining solutions (e.g. meninges, tightly packed axon bundles), which could in turn translate into asymmetrical staining artefacts. To facilitate the identification of specific artefact patterns, we also prepared 87 second to fourth dorsal slices (**Fig. 1a3**) which, while also containing all histological layers, present a symmetrical structure that allows free diffusion of staining solutions through their shortest dimension. Additionally, to account for possible effects only attributable to larger specimen sizes, we prepared 8 and 4 large samples consisting of complete and half olfactory bulbs, respectively, adding up to a total of 427 samples.

**Fig 1.**
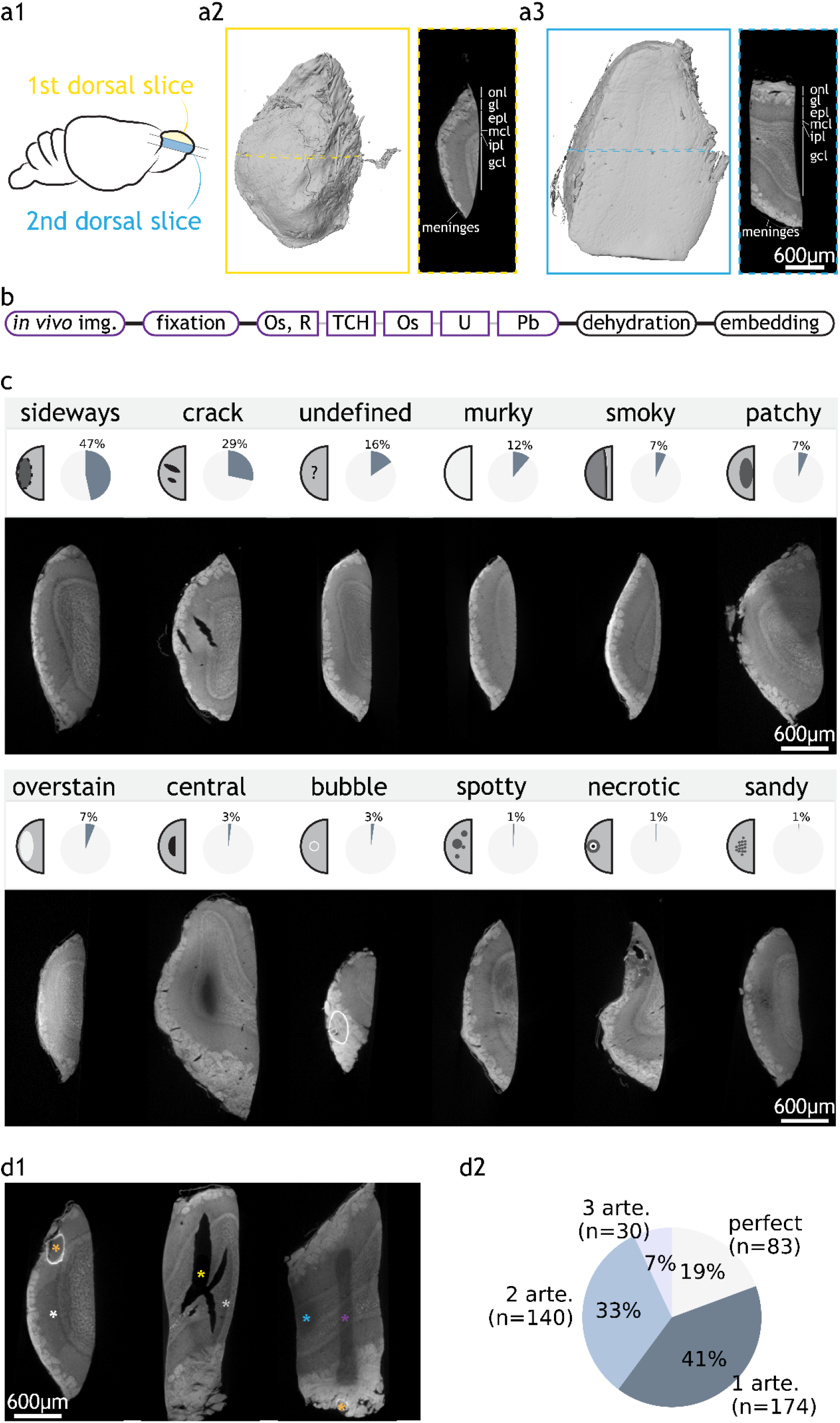
Categorization of staining artefacts from mouse olfactory bulb samples. **(a1)** Positions of the first and second dorsal slices taken from the mouse olfactory bulb. **(a2)** LXRT volume of a first dorsal slice and LXRT image of the dashed region in the volume. **(a3)** same as **(a2)** but for a second dorsal slice. **(b)** Structure of an SPP. Staining is split into key steps. Steps that vary between SPP tests are shown in coloured boxes. **(c)** 12 types of artefacts: name, schematic of appearance, overall occurrence rate in OB samples and LXRT image of a representative sample. **(d1)** Examples of samples with co-occurring artefacts. They respectively contain ‘sideways’ (white asterisk) and ‘bubble’ (orange); ‘crack’ (yellow) and ‘smoky’ (grey); and ‘undefined’ (purple), ‘patchy’ (blue) and ‘bubble’ (orange). **(d2)** Number and percentage of OB samples with zero, one or multiple artefacts. Abbreviations: onl: olfactory nerve layer, gl: glomerular layer, epl: external plexiform layer, mcl: mitral cell layer, ipl: internal plexiform layer, gcl: granule cell layer, img.: imaging. Os: osmium tetroxide, R: reducing agent e.g. potassium ferricyanide, TCH: thiocarbohydrazide, U: uranyl acetate, Pb: lead aspartate, arte.: artefact. Scale bars are shared across all images in the same panel.

We processed the samples following a standard protocol structure (Briggman et al., 2011; Tapia et al., 2012; Wilke et al., 2013; Hua et al., 2015; Mikula and Denk, 2015; Pallotto et al., 2015; Wanner et al., 2016; Genoud et al., 2018) (**Fig. 1b**). After all *in vivo* studies had finished on the animal, the brain tissue was fixed, stained with osmium tetroxide, reduced with potassium ferrocyanide, incubated with the bridging agent thiocarbohydrazide, and further stained with osmium tetroxide, uranyl acetate and lead aspartate. Finally, samples were dehydrated in a series of ethanol solutions, embedded in hard epon resin and cured in an oven for 72h at 60°C. A number of variations were implemented to this protocol backbone with the aim to optimise the yield of samples holding optimal staining quality. Staining quality was routinely assessed with X-ray tomography using a laboratory-based micro-CT (LXRT), a technique well-suited to resolve the different histological layers (Bosch et al., 2021) as well as the dynamics of heavy metal diffusion when staining brain tissue (Ströh et al., 2021).

We observed a wide variety of SPP-derived artefacts (**Fig. 1c**). After examining all stained samples, we described all artefacts into eleven distinct types (**Supp. Table 1**). Each sample was assessed independently for the presence of any of the artefacts. Notably, more than one artefacts can co-occur in one sample (**Fig. 1d**). Some samples (16%, 66/427) showed poor staining but could not be categorised into any of the defined artefacts and were collectively typed as ‘undefined’. Finally, around 20% of the samples do not contain any artefact and were referred to as ‘perfect’ (**Fig. 1d2**). In our samples, the most frequently observed artefacts were ‘sideways’ and ‘crack’, which collectively occurred in 60% of all samples. Other insights could be extracted from the prevalence (% of experiments affected) and penetrance (% of affected samples within an experiment) of the artefacts (**Supp. F1**), yet we focussed our attention on understanding the aetiology of ‘sideways’ and ‘crack’, since even modest improvements in the occurrence of these two major artefacts would convey a meaningful improvement in the overall experimental yield.

The ‘sideways’ artefact consists of a poorly stained region which is fully enclosed within the sample and displaced closer towards the brain surface compared to the sliced surface in first slices (**Fig. 2a**). The transition from well stained to poorly stained region may be abrupt, gradual, or both, giving ‘clear’, ‘fuzzy’, and ‘compound boundary sideways’, respectively (**Fig. 2b**). In second slices, where two sliced surfaces make up most of the surface area, the ‘sideways’ artefact occurs at a different anatomical location yet the boundaries remain closer to the brain surface than to the sliced surfaces (**Fig. 2a**). Moreover, in first slices where the brain surface was locally perturbed (because of e.g. some broken meninges exposing the underlying tissue), the distance from that point in the brain surface to the artefact border was similar to the distance between the artefact border and the sliced surface (**Fig. 2a**). Finally, the overall pattern resembles that of a staining artefact previously described (Hua et al., 2015; Mikula and Denk, 2015) (**Supp. F2**).

**Fig 2.**
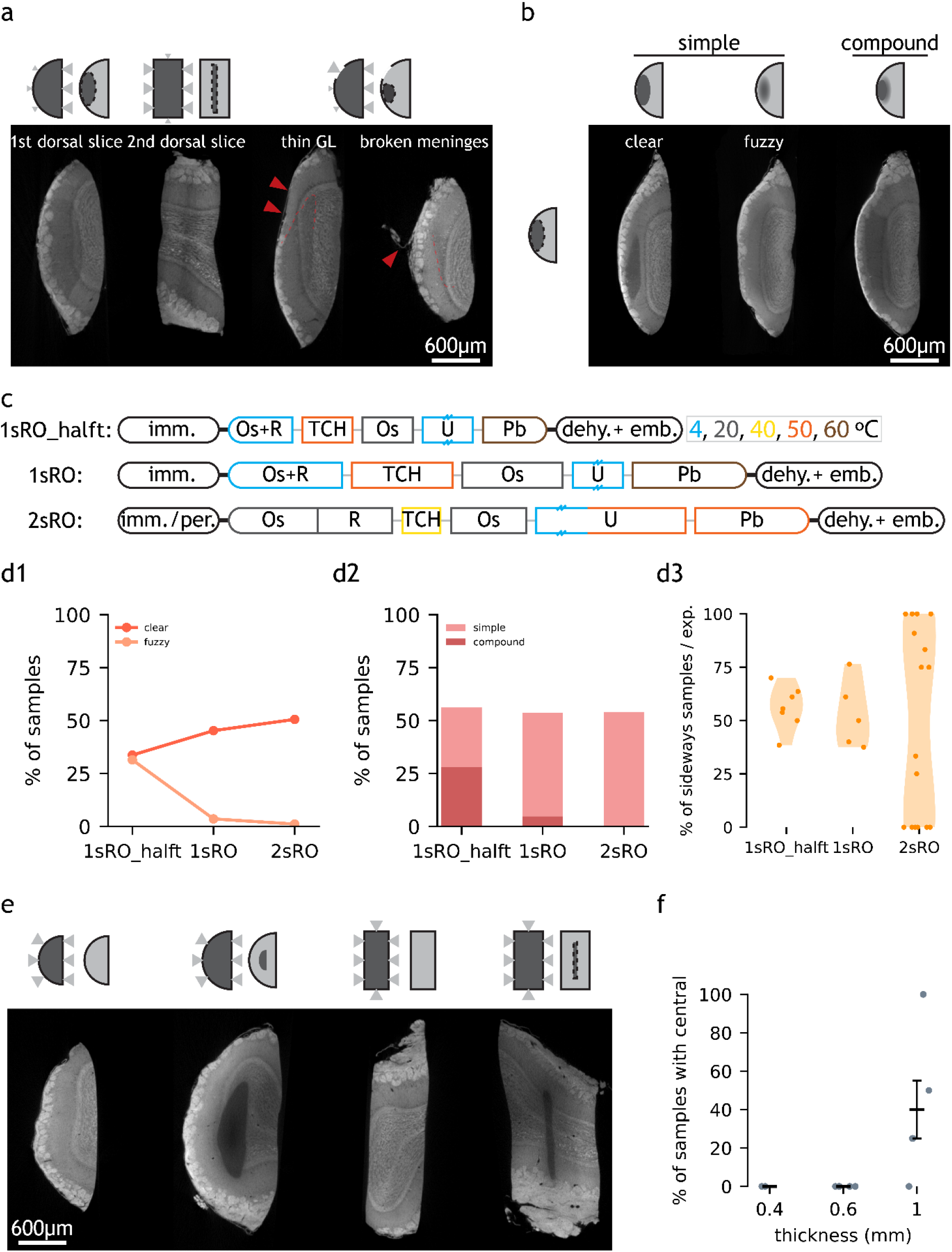
Occurrence and proposed mechanism of ‘sideways’ and ‘central’ artefacts. **(a)** LXRT images of four samples with ‘sideways’ and schematics of hypothesised causes. In the last two samples, the positions of thin GL and broken meninges are highlighted with arrowheads, and part of the ‘sideways’ boundary is marked with a dashed line. In schematics, the size of triangles represent the diffusion rate. **(b)** Subdivision of ‘sideways’ by boundary type and by complexity. **(c)** Summary of three SPPs. Staining is split into key steps. Length and colour of staining steps represent incubation time and temperature, respectively. Double dash on box represents overnight incubation. Thin lines connecting staining reagents represent water washes. **(d1)** Occurrence rate of ‘sideways’ with ‘clear’ and ‘fuzzy’ boundaries in the three SPPs. **(d2)** Occurrence rate of ‘simple’ or ‘compound’ ‘sideways’ in the three SPPs. **(d3)** Penetrance of ‘sideways’ in experiments processed with the three SPPs. Each dot represents one experiment. **(e)** Samples with (2nd and 4th, 1 mm-thick) and without (1st and 3rd, 0.6 mm-thick) the ‘central’ artefact. **(f)** Occurrence of ‘central’ depends on slice nominal thickness. Each data point represents the % of samples with ‘central’ artefact within an experimental batch. Error bar shows mean +/- standard error of mean. Abbreviations: GL: glomerular layer, imm. : immersion, per.: perfusion, dehy.: dehydration, emb,: embedding, Os: osmium tetroxide, R: potassium ferricyanide, TCH: thiocarbohydrazide, U: uranyl acetate, Pb: lead aspartate. Scale bars are shared across all images in the same panel.

Heavy metals passively diffuse through soft tissue following a quadratic kinetic (Mikula and Denk, 2015; Ströh et al., 2021), i.e. staining three times the depth takes nine times longer. From our observations, we hypothesised that the brain surface is less permeable than the sliced surface to the diffusion of heavy metals. If so, reagents would reach locations through the sliced (permeable) border more easily. This combined with insufficient incubation times could lead to the ‘sideways’ artefacts we observed.

In order to assess whether diffusion time affected the presence of this artefact, we modified the protocol so the duration of all staining steps was doubled (**Fig. 2c**). This led to a slight change in the pattern of the ‘sideways’ artefact - with it now displaying only clear boundaries (**Fig. 2d1-d2**).

A second mechanism that could contribute to the ‘sideways’ artefact is the reduction of osmium VIII to osmium IV inside the tissue during staining (Hua et al., 2015). A prior study showed mm-thick mouse brain tissue samples could be evenly stained for SBEM imaging if the reducing agent was presented after the tissue had been already stained with osmium tetroxide (instead of presenting the fresh tissue with reduced osmium tetroxide). We refer to this protocol as ‘2-step reduced osmium’ (2sRO) (**Fig. 2c**). 2sRO also differed in that the incubation of the bridging agent TCH was milder (in time and in temperature), the uranyl acetate incubation (typically overnight at 4°C) was extended with a short (2h) hot (50 °C) period and the incubation of lead aspartate was milder (in temperature). Finally, this protocol employed a distinct fixation method - perfusion - to the one used before - immersion. We implemented the complete 2sRO to replicate its reported performance and compared it across our benchmarked protocol (1sRO). Indeed, experiments stained with 2sRO displayed a distinct penetrance of the ‘sideways’ artefact, which was entirely absent in all samples of 38% of the experiments (6/16) (**Fig. 2d3**).

‘Sideways’ is not the only diffusion-related artefact we observed. ‘Central’ displays a poorly stained region fully enclosed inside the sample, but in this case the unstained region presents virtually no metal staining - with no histological structure revealed by LXRT, and its location appears equally distant to all sample borders (**Fig. 2e**). It is similar to previously reported artefacts associated with insufficient staining times (Hua et al., 2015; Mikula and Denk, 2015; Ströh et al., 2021), and it was completely absent in slices with thickness smaller or equal than 0.6 mm (**Fig. 2f**).

Altogether, we identified two possible mechanisms behind the most prevalent artefact ‘sideways’: diffusion kinetics of the staining solutions and reduction of osmium tetroxide during the first staining step, and the implemented changes removed this artefact from all samples in 38% of the experiments.

The second most prevalent artefact was ‘crack’ (**Fig. 1c**), whose presence precludes any further analysis on the affected sample: if this artefact occurs in the volume of interest, samples become unusable as cellular features cannot be annotated across even small cracks. In fact, even if ‘crack’ artefacts were found outside of the volume of interest, the sample should also be treated as unusable because smaller cracks were likely to be found across the rest of the volume. We further classified the observed cracks into two non-mutually-exclusive subtypes, based on whether the crack did not follow any histological feature (‘crack I’) or whether it followed histological borders (‘crack II’) (**Fig. 3a**).

**Fig 3.**
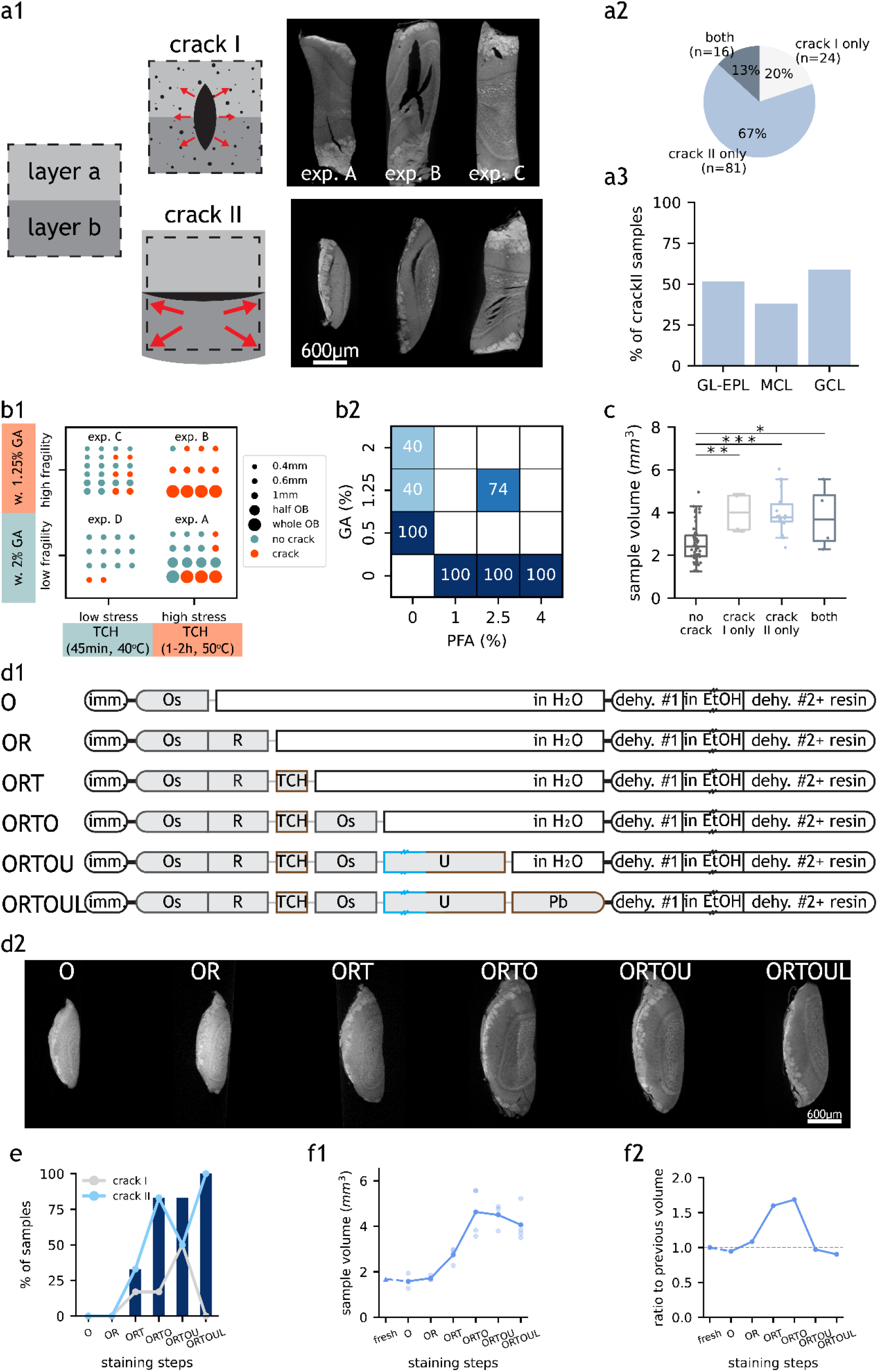
Occurrence and proposed mechanisms of ‘crack’ subtypes. **(a1)** LXRT images of samples with ‘crack’ subtypes and schematics of their hypothesised causes. In schematics, ‘layer a’ and ‘layer b’ represent two tissue layers and the dashed box represents the starting tissue volume i.e. when fresh. Dots represent gas bubbles. Arrows represent expansive forces experienced by the tissue. Top, 0.6 mm-thick representative samples from experiments A, B and C referred to in (b1). Bottom, samples showing ‘crack II’ at GL-EPL boundary, MCL, and GCL, respectively. **(a2)** Number and percentage of ‘crack’ samples with only ‘crack I’ or ‘crack II’ or with both ‘crack’ subtypes. **(a3)** Percentage of ‘crack II’ samples with the artefact occurring at GL-EPL boundary, MCL or GCL. **(b1)** ‘crack’ occurrence in samples from four experiments with different TCH incubation conditions and GA concentrations. **(b2)** ‘crack’ occurrence in 0.6 mm-thick samples fixed with different fixatives. **(c)** Volume of samples that do and do not contain ‘crack’. Each dot represents one sample. The boxes extend from the first to the third quartile, with the line representing the median. The whiskers extend to the most extreme, non-outlier data points. (*: p<0.5, **: p<0.005, ***: p<0.0005, two-tailed paired t-test). **(d1)** Experimental design of a step-by-step investigation of artefact occurrence. Double dash on box represents overnight incubation. Length and colour of staining steps represent incubation time and temperature, respectively. **(d2)** LXRT images of representative samples from each experimental group described in (d1). **(e)** Occurrence of ‘crack I’ (grey line), ‘crack II’ (blue line) and ‘crack’ (blue bars) in each experimental group described in (d1). **(f1)** Estimated fresh sample volume (triangle) and measured final sample volume (dots) for samples from experimental groups described in (d1). Each light dot represents one sample. Each dark dot represents the average of all samples from an experimental group. A dashed line connects estimated and measured volumes and solid lines connect the measured sample volumes. **(f2)** Ratio of the mean sample volume of each experimental group to that of the previous group. A dashed line marks ratio = 1 i.e. no volume change. Abbreviations: GL: glomerular layer, EPL: external plexiform layer, MCL: mitral cell layer, GCL: granule cell layer, w.: with, GA: glutaraldehyde, TCH: thiocarbohydrazide, PFA: paraformaldehyde, imm. : immersion, Os: osmium tetroxide, R: potassium ferricyanide, TCH: thiocarbohydrazide, U: uranyl acetate, Pb: lead aspartate, dehy.: dehydration, EtOH: ethanol. Scale bars are shared across all images in the same panel.

‘Crack’ might be generated by expansive forces overwhelming the forces that keep the tissue together (**Fig. 3a1**). Therefore, we expected chemical fixation to play a role in preventing the appearance of cracks - by making the tissue less fragile. Paraformaldehyde and glutaraldehyde are the two most commonly used fixatives for EM, employed either alone or in combination (Morris, 1965). Glutaraldehyde provides a stronger fixation, derived from its two aldehyde groups that enable reactions cross-linking protein molecules (Kiernan, 2000). We compared sample batches fixed with commonly used fixative solutions employing distinct concentrations of glutaraldehyde. On the other hand, it has been reported that the bridging agent TCH generates gaseous nitrogen as a subproduct when binding to the fixed osmium (Guha and De, 1924; Seligman et al., 1966; Mikula and Denk, 2015).

We assumed the stress generated would be related to the amount of gas created, which in turn should correlate with the amount of TCH-osmium reactions. We thereby devised two stress scenarios, differing in the duration and temperature of the TCH step. We observed that fewer samples carried ‘crack’ artefact when processed with the milder TCH incubation, even when fixed with the milder fixative solution (**Fig. 3b1**), and that crack severity decreased at the same time (**Fig. 3a1**). Moreover, glutaraldehyde was required in the fixative solution in order to prevent the appearance of cracks (**Fig. 3b2**). Finally, cracks of all types were more frequently present in larger samples (**Fig. 3c**).

We then aimed to identify at which point in the staining protocol ‘crack’ appeared by stopping the staining process at every single step (**Fig. 3d**). As expected, ‘crack’ started appearing during the TCH step, and their presence increased progressively throughout the consecutive staining steps (**Fig. 3e**). Moreover, this experiment showed that samples expand during the staining protocol (**Fig. 3d2**), that expansion starts during the TCH step and that samples continue to expand during the following osmium tetroxide step (**Fig. 3f**).

Altogether, ‘crack’ artefact can be prevented by including glutaraldehyde in the fixative and by minimising the TCH reaction.

Next, we evaluated the effect of prior *in vivo* 2P imaging on the experimental yield of the SPPs (**Fig. 4**). The throughput (the number of samples processed per batch) was severely reduced by the addition of the *in vivo* imaging step (**Fig. 4a**). Nevertheless, on average the success rate of the experiment remained unaltered by the incorporation of this imaging modality (**Fig. 4b1**), despite introducing some 2P-specific artefacts (**Fig. 4b3-4**). Edema is considered a source of surgery-derived histology-scale damage (Betz and Coester, 1990; Judkewitz et al., 2009). To minimise this factor, we introduced a series of edema-preventing measures before and during surgery: mice were administered with the anti-inflammatory drug dexamethasone before the experiment, ice-cold buffer was frequently applied to cool down the drilling of the skull and laser exposure during imaging was minimised whenever possible. Nevertheless, we did not observe a significant effect of these additional measures (**Fig. 4c**).

**Fig 4.**
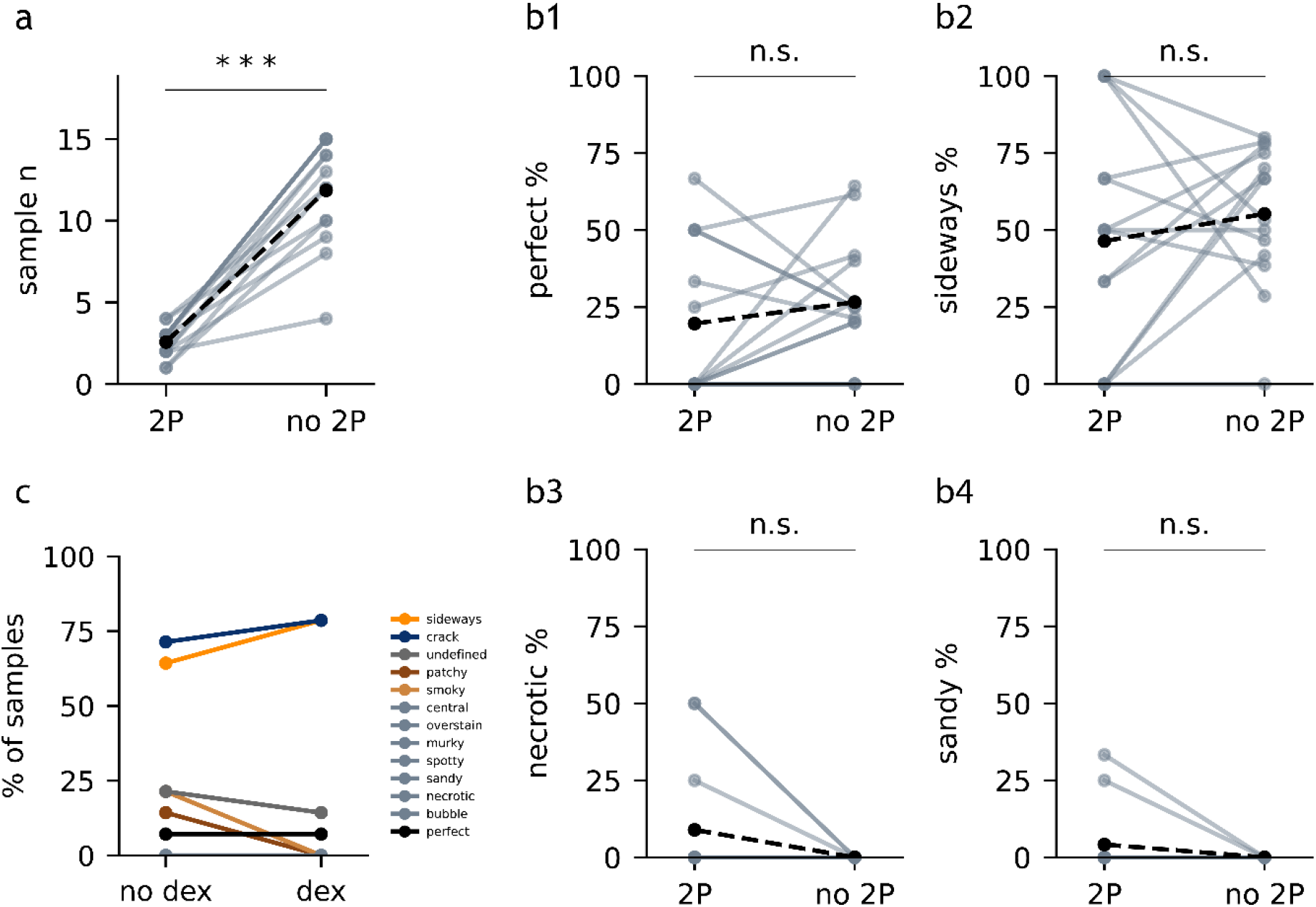
The effect of 2P and dexamethasone on artefact occurrence. **(a)** Number of samples with (‘2P’) and without *in vivo* functional imaging (‘no 2P’) from each experiment. Each light dot represents ‘2P’ samples or ‘no 2P’ samples of one experiment. Solid lines connect ‘2P’ samples to ‘no 2P’ samples from the same experiment. Black dots represent the mean. Dashed lines connect the mean values. Asterisks represent statistical significance (***: p<0.0005, two-tailed paired t-test). **(b1)** Percentage of perfect samples from each experiment. **(b2-4)** Percentage of samples with ‘sideways’ (b2), ‘necrotic’ (b3) and ‘sandy’ (b4). **(c)** Artefacts in samples administered with and without dexamethasone within the same experiment (n = 13 and 14, respectively).

In sum, we optimised the SPP to obtain good quality images from all modalities with sufficient yield. At this point we turned our attention towards the second main factor affecting the throughput of the correlative CMI workflow: the quality of the registration of the different modalities into a multimodal map.

In order to reliably identify neurons functionally-recorded *in vivo* in the anatomical map acquired *ex vivo*, the datasets arising from each imaging modality need to be spatially correlated at cellular precision (Helmstaedter, 2013). By identifying conserved pairs of landmarks in two modalities reporting the same space, a warp field relating the two spaces can be computed (**Fig. 5a1**) (Bogovic et al., 2016). This set of landmarks was termed ‘warping landmarks’. This warp field can later be used to translate any spatial annotation across the two coordinate systems (**Fig. 5a2**). Blood vessels can be used to set warping landmarks for correlating *in vivo* 2P and SBEM datasets, since their structures can be reliably retrieved by both modalities. They often describe chiral spatial patterns that occur at a density potentially sufficient to map tissue at single cell body precision (Bock et al., 2011; Briggman et al., 2011; Lee et al., 2016; Bosch et al., 2020; Bosch et al., 2021; Miettinen et al., 2021). As blood vessels are also resolved by SXRT, they can be used as a source for landmarks for warping across the three dataset modalities (**Fig. 5b**).

**Fig 5.**
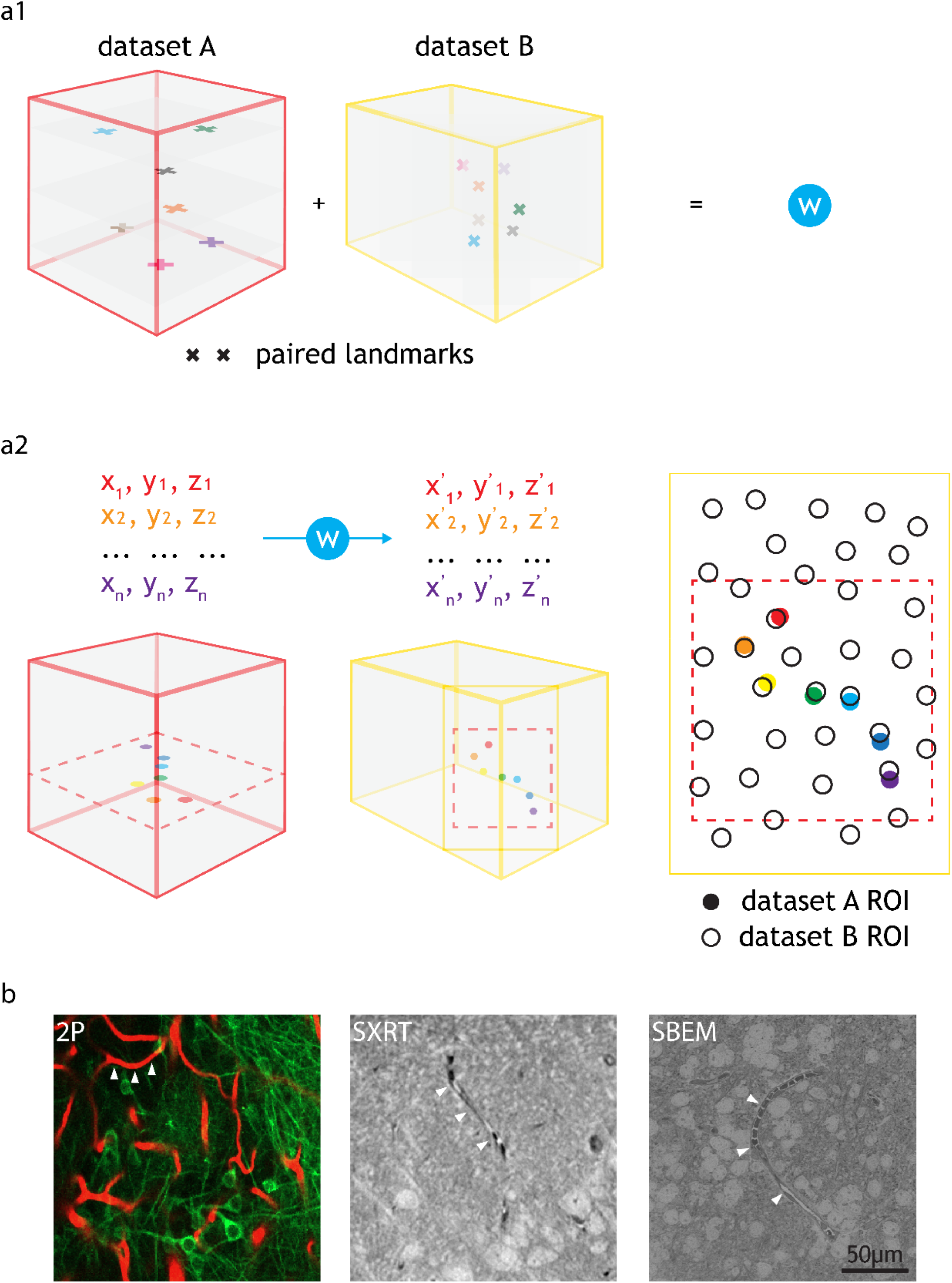
Warping using common features e.g. blood vessels and cell correlation by annotation warping. **(a1)** Common features found throughout the volume of two datasets (i.e. paired landmarks) are used to calculate a warping function (‘W‘). **(a2)** The warping function can be used to warp any annotations from one dataset to another. The inputs needed are the x, y, z coordinates of the annotations in the native dataset space and the outputs are the x, y, z coordinates of those annotations in the target dataset space. Annotation warping is used to correlate ROIs from two datasets. ROIs can be correlated as long as the warped ROI falls to a position that is closer to the target ROI than any other ROI. For (a1-a2), cubes represent datasets (e.g. 2P, LXRT, SXRT, SBEM), acquired from the same piece of tissue. The datasets may differ in the dimensions, voxel sizes and the exact region of the sample, differences here represented by cube dimensions, landmark sizes and landmark positions, respectively. Other differences may involve a reversed order in one of the spatial dimensions (such as an horizontal flip in all images), making it necessary to use chiral structures as a source for warping landmarks. **(b)** Example images showing blood vessels (arrowheads) resolved by 2P, SXRT and EM datasets.

We aimed to quantify the precision at which one could warp datasets obtained by those three modalities in a previously established correlative workflow (**Fig. 6a-b**) (Bosch et al., 2021). For this purpose, we manually traced blood vessels in the same region spanning (160 µm)^3^ in each original dataset (**Fig. 6a**). Then, these tracings were warped into a common reference space using the warp field calculated with the previously described set of ‘warping landmarks’ (**Fig. 6c**). Blood vessel branching points are unique junctions and many of them (n=39) could be identified across all imaging modalities, defining blood vessel segments (n = 35). We interpolated a defined number of points within each blood vessel segment in all modalities and paired the resulting nodes (**Fig. 6d**). This allowed for calculating the distance across modalities between paired nodes of the blood vessel (**Fig. 6e**). Warping accuracy depends on the accuracy at which landmarks are resolved across the dataset and therefore its value can vary throughout the dataset’s topology. We observed that the range of warping accuracies recorded depended on the pair of imaging modalities being warped: it ranged between 0.6 and 27.3 µm when warping 2P to EM or to SXRT and between 0.2 and 9.9 µm when warping SBEM to SXRT (**Fig. 6e, f**). On average, any point in the blood vessel could be mapped at ∼14 µm accuracy between the 2P and the SBEM dataset and at ∼9 µm accuracy between the 2P and the SXRT dataset, and regions between SXRT and SBEM could be mapped at an accuracy of ∼2 µm (**Fig. 6f**,**g**), implying that any region can be mapped across any of the three imaging modalities at single cell precision (**Fig. 6h**).

**Fig 6.**
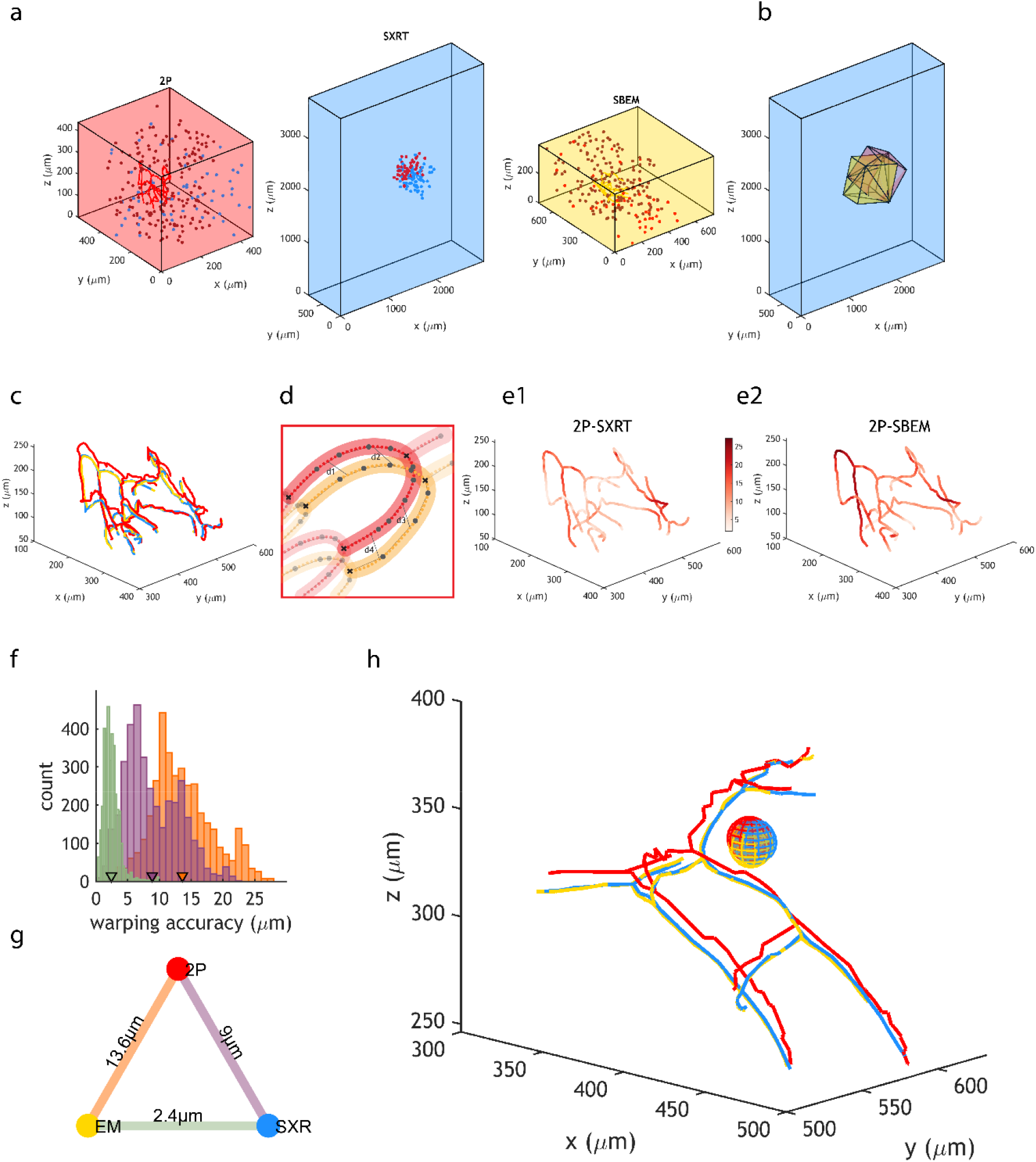
Accurate warping allows for cell correlation among 2P, SXRT and SBEM. **(a)** Position of landmarks used to generate warping functions between two datasets and position of blood vessel tracings used to quantify warping accuracy in 2P, SXRT and SBEM datasets of a sample. Blue, brown and red dots represent landmarks used between 2P and SXRT datasets, between 2P and SBEM datasets and between SXRT and SBEM datasets, respectively. **(b)** Fields of view of 2P (red, warped), SBEM (yellow, warped) and SXRT (blue, native space) datasets. **(c)** Same blood vessels traced in the 2P (red, warped), SXRT (blue, warped) and SBEM (yellow, native space) datasets. **(d)** Quantification of warping accuracy. The nodes in the blood vessel traced in one imaging modality (red) are paired to the equivalent nodes in the other modality (yellow). The distance between two paired nodes (e.g. ‘d1’) reflects the warping accuracy at that location. Large dots represent nodes in the original tracing. Lines represent paths of the original tracing. Small dots represent nodes interpolated along the tracing path. **(e1-2)** Native SBEM blood vessel tracing coloured by the mapped 2P-SXRT and 2P-SBEM warping accuracy, respectively. **(f)** Distribution of warping accuracies (i.e. distances between paired nodes). Purple, orange and green represent 2P-SXRT, 2P-SBEM and SXRT-SBEM accuracies, respectively. Mean warping accuracies are marked by triangles. **(g)** Mean warping accuracy between each pair of imaging modalities. **(h)** Example mitral cell and surrounding blood vessels warped from 2P (red) and SXRT (blue) datasets into SBEM (yellow) space. The diameter of the sphere is 20um, similar to the diameter of a mitral cell soma.

## Discussion

We have identified and addressed two major bottlenecks in the performance of a correlative multimodal imaging pipeline capable of resolving the structure and function of a neural circuit in the mouse olfactory bulb: the yield of sample preparation and the accuracy of dataset correlation across 2P, SXRT and SBEM modalities.

Our study focuses on mouse olfactory bulb (OB) dorsal slices. The OB contains the first neuronal circuit in the olfactory sensory pathway (Mori et al., 1999), the glomerular column. The architecture of these columns is compact (<1 mm^3^) and modular (>1000 columns in each OB hemisphere) (Schwarz et al., 2018) and their anatomical location makes them accessible for *in vivo* studies involving minimally invasive surgeries (e.g. imaging intracellular [Ca^2+^] through a cranial window). Combined, these properties make the OB a good model for studying systems neuroscience in the mammalian brain (Schaefer and Margrie, 2007). For *in vivo* experiments aimed at recording the circuit’s function, 2P datasets can map a dorsal field of view of ∼mm^2^, but depth resolution of 2P is poorer than the lateral one (Chaigneau et al., 2011). Moreover, its signal-to-noise ratio decreases with the depth of the region being imaged, making it increasingly challenging to retrieve a signal beyond a depth of 0.5 mm deep inside the brain (Wachowiak et al., 2013; Ackels et al., 2020).

Our horizontal slices provided a maximal overlap between the 2P-imaged region and the stained tissue that would later be imaged at higher structural resolution with SXRT and SBEM. However, these dorsal sections present anatomical asymmetries: one of their sides is a vibratome-cut surface, whereas the other side is the anatomically-preserved surface of the brain. We generated a set of samples that would present a more symmetrical disposition to diffusing solutions: second slices from the same brain region.

Two main artefacts dominated the sample preparation protocol (SPP). An asymmetrical unstained region here denoted ‘sideways’ was the most frequent artefact type. Its pattern was reminiscent of an artefact defined previously (Hua et al., 2015; Mikula and Denk, 2015) although in our samples it displayed an asymmetrical morphology (**Supp. F2**). We found it to be related to two independent parameters: the diffusion of heavy metals through the tissue and the reduction of the first osmium tetroxide staining step. This is consistent with its symmetrical appearance in previous reports that analysed punch-dissected tissue cylinders presenting a symmetrical structure to freely diffusing solutions (Hua et al., 2015). Addressing those two factors allowed for removing its presence in all samples in a subset of experimental batches.

The next most common artefact was the presence of cracks in the stained tissue, possibly generated by an imbalance between expansive and cohesive forces in the tissue during sample preparation. It is known that osmium tetroxide can provoke both the expansion and the shrinkage of brain tissue samples during staining (Ströh et al., 2021) and that the bridging agent TCH generates gaseous nitrogen as a by-product of binding to fixed osmium (Guha and De, 1924; Seligman et al., 1966; Mikula and Denk, 2015). Consistent with that, we found that a milder TCH reaction can minimise the generation of cracks in samples extending multiple mm^3^ in volume, while not compromising the overall staining quality. On the other hand, strong sample fixation with glutaraldehyde was necessary to prevent tissue from cracking during the staining process. Finally, larger samples displayed cracks more frequently. Altogether, these findings allowed extracting simple specific requirements for future protocols to minimise the appearance of cracks: (1) employing fixatives containing at least 1.25% glutaraldehyde, (2) reducing the strength of the TCH reaction, and (3) minimising the smallest dimension of the sample being processed whenever possible. Further, albeit more difficult to implement improvements are: (1) finding optimal formulations for fixative solutions for a specific sample, (2) replacing TCH for a reagent that does not create bubbles - such as pyrogallol (Mikula and Denk, 2015), and (3) minimising the sample expansion processes by e.g. counteracting osmium-derived effects with suitable additives (Hua et al., 2015; Ströh et al., 2021).

A subset of the experiments reported were obtained by chemically fixing the tissue by immersion in fixative (**Supp. Table 2**). All artefacts in immersion-fixed samples could be diagnosed by observing an LXRT dataset covering the entire sample. Such 3D datasets can be obtained at a rate of multiple samples per day with benchtop LXRTs (Metscher, 2009; Bushong et al., 2015; Gutiérrez et al., 2018) and of nearly one sample per minute at specialised hard X-ray microtomography beamlines at synchrotron X-ray facilities (Walker et al., 2014; Rau, 2017; Harkiolaki et al., 2018; Bosch et al., 2021; Walsh et al., 2021). A distinct approach could involve fixing the samples via intracardiac perfusion of the fixative (Au - Gage et al., 2012) (**Supp. F3**). Doing so could prevent some artefacts that might be specific to immersion-fixation (**Supp. F3a**) and would remove the requirement of a fast dissection of the tissue of interest, making this approach compatible with processing some regions that might be out of reach or processing multiple different regions in the same animal. On the other hand, maintaining ultrastructural integrity is generally more challenging with perfusion (Palay et al., 1962; Crang et al., 1988; Maunsbach and Afzelius, 1999; Tao-Cheng et al., 2007; Glauert and Lewis, 2014) (**Supp. F3b**). It is worth noting that evaluating sample quality by its ultrastructure has a lower throughput than relying on LXRT for that purpose. Therefore, choosing perfusion as a fixation method could create a quality control bottleneck in the experimental pipeline (**Supp. F3c**).

The yield of perfectly stained samples in an experiment was not altered when extending the pipeline with prior *in vivo* 2P imaging. However, those experiments have a much lower throughput (**Fig. 3a**). A low throughput combined with a moderate yield can make it challenging to predict future stock of samples with both optimal 2P imaging and optimal staining. Importantly, sample preparation does not end with staining: trimming the specimen will be necessary to accommodate its dimensions to some acquisition techniques (**Supp. F3c**) (Karreman et al., 2016; Bosch et al., 2021; Ronchi et al., 2021), hard X-ray tomography imaging may inflict damage from radiation dose and temperature changes (Henderson, 1995; Howells et al., 2009; Du and Jacobsen, 2018), volume electron microscopy techniques imply slicing or destruction of the sample and therefore add their own risk of a specific sample becoming irreversibly damaged, and optimal targeting of the features of interest to be imaged with the distinct modalities precludes extracting the information sought by the initial experimental question (Bosch et al., 2020; Walter et al., 2020). In light of these successive challenges, it is imperative for a successful CMI experiment to prepare a sufficient number of samples at the first stage. Given the low throughput nature of *in vivo* 2P experiments (in particular in e.g. awake or even behaving animals), this can currently partially be achieved by e.g. obtaining samples from multiple brain regions in a given experiment or by coordinated efforts of multiple researchers or even research labs. Notably, any improvement to the SPP will reduce this initial experimental burden and allow for increasingly complex physiological analysis to be combined with structural investigations.

Finally, we demonstrated that tissue locations imaged *in vivo* can be recalled at single-cell precision in an SXRT dataset. Because SXRT datasets can be obtained on multi-mm^3^ specimens, this strategic improvement provides an early readout of circuit structure and can inform on how to precisely trim precious specimen, preserving virtually all 2P-imaged cells in a trimmed specimen for follow-up higher resolution structural imaging techniques that impose limitations on sample dimensions and require extraordinary sample quality control standards. Direct 2P-SXRT correlation therefore generates experimental compatibility of *in vivo* 2P with not only the different implementations of volume EM but also with other synchrotron hard X-ray modalities such as nanoholotomography (Kuan et al., 2020), laminography (Helfen et al., 2013; Witte et al., 2020) or ptychography (Rodenburg et al., 2007; Holler et al., 2017; Shahmoradian et al., 2017). Adding non-destructive techniques to the workflow not only enables more diverse insights to be recalled from the same sample, but also makes it more resilient ensuring insights are obtained from a larger yield of samples. Altogether, we are confident that improved staining and reliable warping will enable exhaustive studies retrieving physiology, circuit structure and synaptic signatures in targeted regions of slices covering multiple mm^3^.

Our optimisation was aimed at obtaining a single sample preparation protocol that is compatible with both SXRT and SBEM, since staining for the latter did provide a sufficient signal-to-noise ratio in the former. However, it is possible that better SXRT datasets could be obtained if the samples were prepared following specially SXRT-optimised preparation protocols. Those could, for example, aim to maximise the phase contrast in the detected X-ray beam while minimising the absorption and thereby bring benefits in terms of both image contrast and specimen size. Doing so could be envisaged by using weaker metal stainings. This could modify, in turn, the landscape of compatible techniques for future multimodal imaging approaches in systems neuroscience and biological soft tissue research more generally, e.g. enable to extract insights on molecular composition and transcriptomic expression in the tissue alongside its function and structure (Ståhl Patrik et al., 2016; Domart et al., 2020; Close et al., 2021).

Altogether, we report improvements in two limiting steps of a correlative multimodal imaging pipeline for systems neuroscience in the mouse olfactory bulb in sample preparation and in dataset correlation accuracy. These insights will allow us to make informed decisions to troubleshoot and improve the yield of the existing protocols and to design novel pipelines relying solely on hard X-rays to retrieve the location of the cell bodies of interest. These frameworks will pave the way for the design of more resilient workflows to interrogate both structure and function of neural circuits in the mammalian brain.

## Acknowledgements

We are grateful to the biological research and electron microscopy science technology platforms of the Francis Crick Institute. We thank Marta Pallotto, Kara Fulton and Kevin Briggman for support and critical discussions on the initial configuration of the staining protocol, to Manuel Berning for support in developing the warpAnnotations toolbox, to Norman Rzepka for support in the webknossos infrastructure, and to the members of the sample preparation working group of the volumeEM community for insightful discussions on the topic.

## Data availability statement

All data tables and scripts used for the artefact analysis, links to LXRT datasets of representative samples of all artefact categories and links to correlated blood vessel annotations on the 2P, SXRT and SBEM datasets are available in the shared repository **protocolBLAST** (https://github.com/FrancisCrickInstitute/protocolBLAST).

The warping landmarks, functions and a toolbox to analyse webKnossos annotations are available in the shared repository **warpAnnotations** (https://github.com/FrancisCrickInstitute/warpAnnotations).

## Ethics statement

All animal protocols were approved by the Ethics Committee of the board of the Francis Crick Institute and the United Kingdom Home Office under the Animals (Scientific Procedures) Act 1986.

## Author contributions

YZ, ATS and CB conceptualised the research project and designed experiments; YZ, TA, AP, MCZ, AB and CB performed experiments and acquired data; YZ, ATS and CB analysed and interpreted the data; YZ, ATS and CB wrote the manuscript draft; YZ and CB generated the figures, and all authors reviewed and provided input to the manuscript.

## Supplementary Material

The Supplementary Material for this article is available online at the same location as the manuscript.

