## Supplementary Figures 1-4 for "Sample preparation and warping accuracy for correlative multimodal imaging in the mouse olfactory bulb using 2-photon, synchrotron X-ray and volume electron microscopy"

### Supplementary Figures and Tables

a

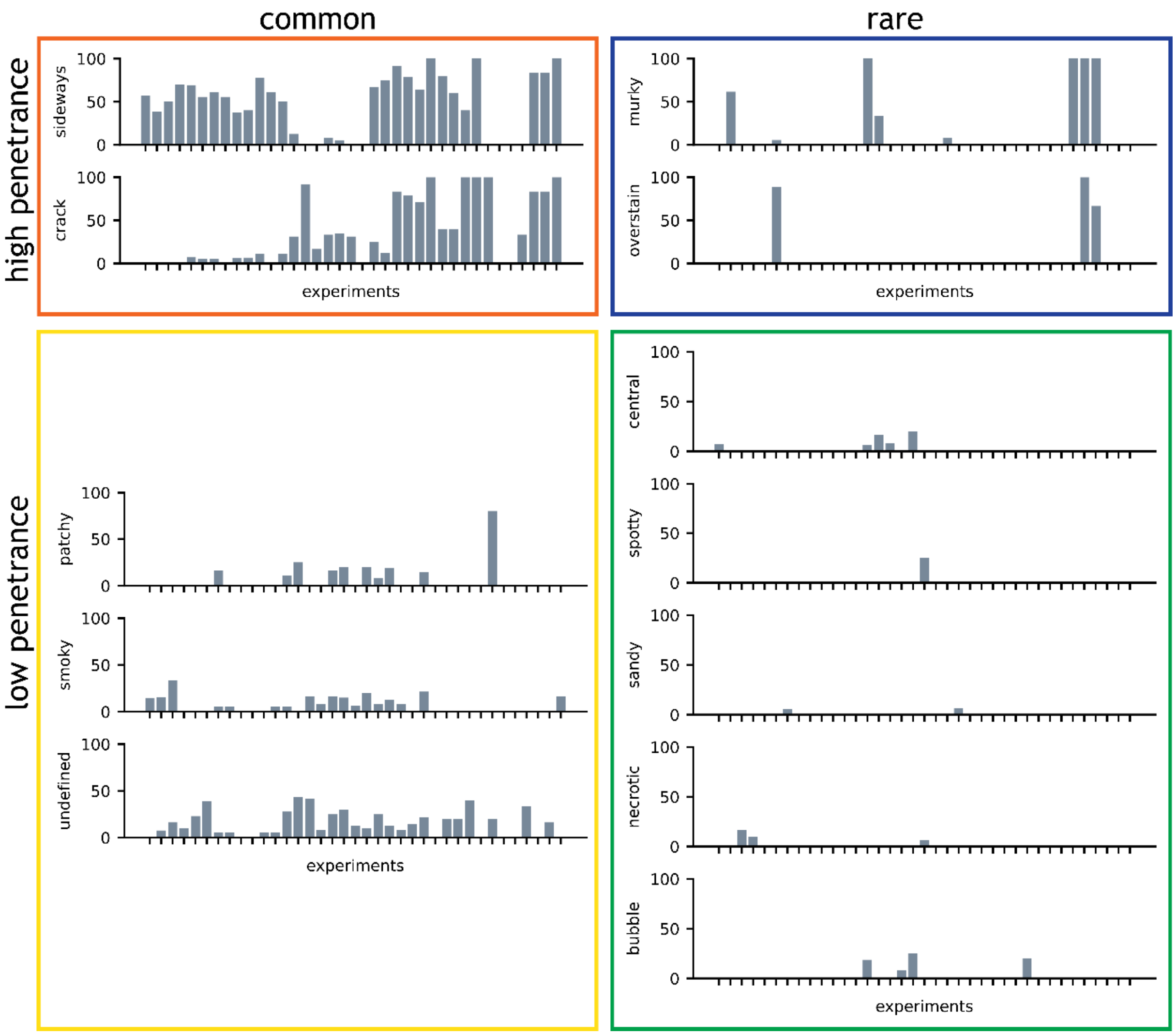


**Supp. F1. Prevalence and penetrance of artefacts.**

**(a)** Artefacts grouped by prevalence and penetrance. Prevalence describes the number of experiments containing samples with an artefact. Penetrance describes the percentage of samples affected by an artefact within one experiment.

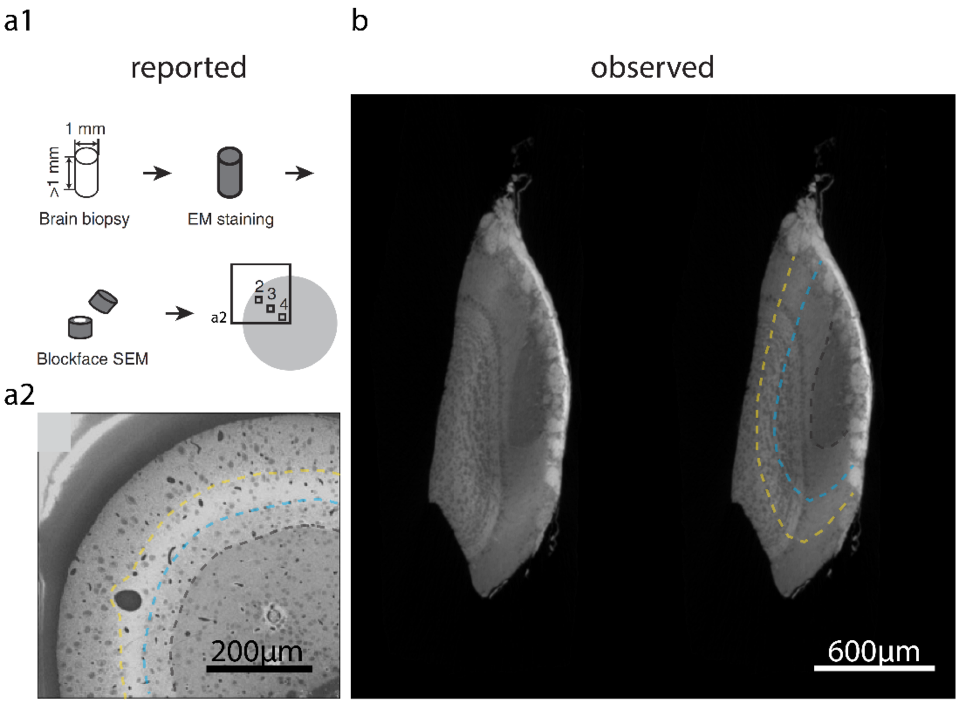


**Supp. F2. Resemblance of ‘sideways’ to previously reported artefacts.**
**(a)** Staining artefact previously described, displaying multiple symmetrical staining abnormalities. **(a1)** Samples were punch-dissected cylinders of adult mouse brain tissue. **(a2)** SBEM image of a cross-section of the cylinder. The boundaries of the different staining abnormalities are marked with blue and yellow dashed lines. **(b)** LXRT images of a first dorsal OB slice sample from this study containing ‘sideways’ artefact with ‘compound’ boundary. The boundaries of staining abnormalities equivalent to those reported in (a2) are delineated with blue and yellow dashed lines, respectively. Panels **(a1-2)** were adapted from Hua et al. (2015).

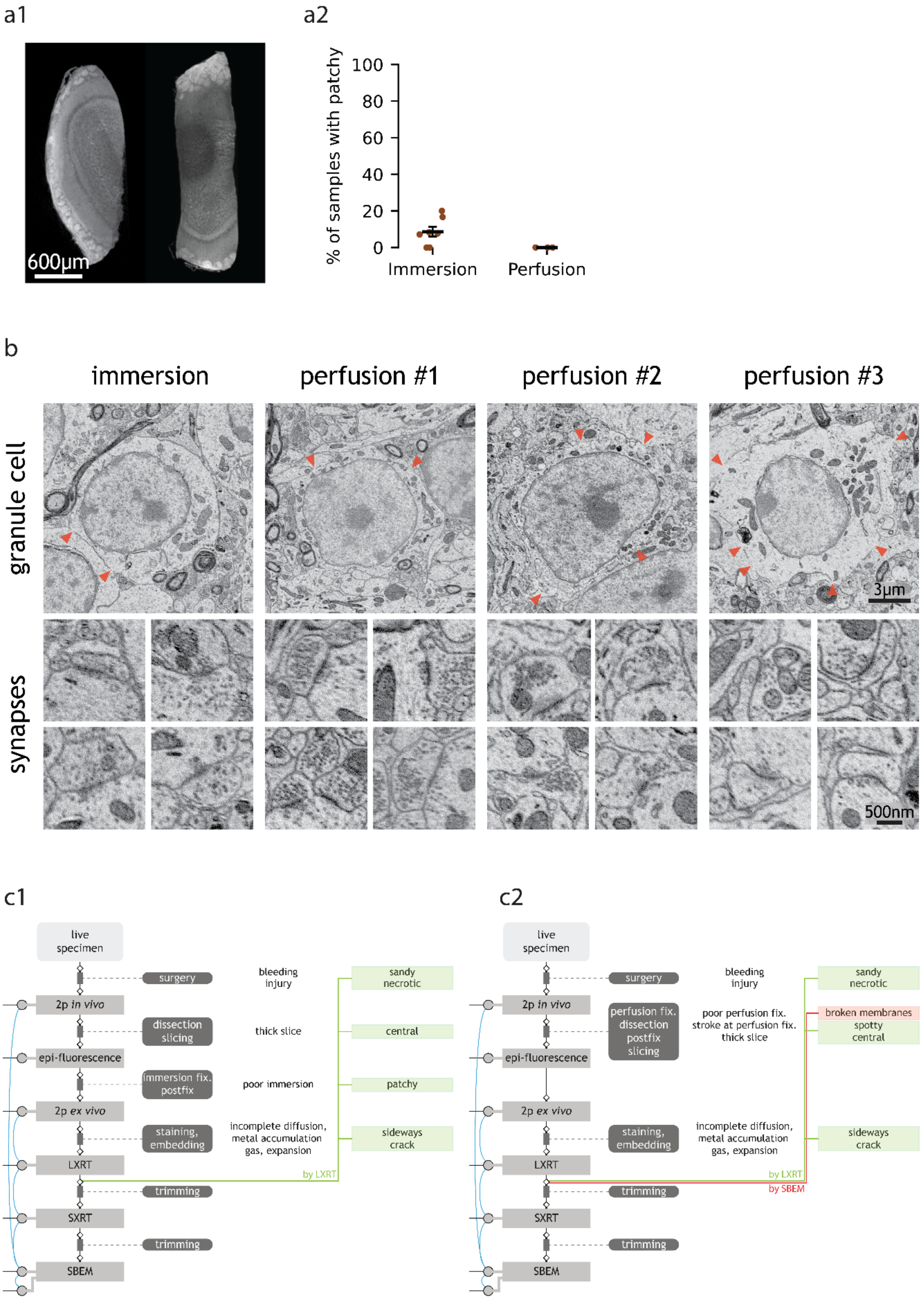

a1
600μm
a2
% of samples with patchy
100
80
60
40
20
0
Immersion
Perfusion
b
immersion
perfusion #1
perfusion #2
perfusion #3
granule cell
synapses
3μm
500nm
c1
live specimen
2p in vivo
epi-fluorescence
2p ex vivo
LXRT
SXRT
SBEM
surgery
dissection slicing
immersion fix. postfix
staining, embedding
trimming
trimming
bleeding injury
thick slice
poor immersion
incomplete diffusion, metal accumulation gas, expansion
sandy necrotic
central
patchy
sideways crack
by LXRT
c2
live specimen
2p in vivo
epi-fluorescence
2p ex vivo
LXRT
SXRT
SBEM
surgery
perfusion fix. dissection postfix slicing
staining, embedding
trimming
trimming
bleeding injury
poor perfusion fix. stroke at perfusion fix. thick slice
incomplete diffusion, metal accumulation gas, expansion
sandy necrotic
broken membranes
spotty central
sideways crack
by LXRT
by SBEM

**Supp. F3. Workflows with different fixation methods exhibit different artefacts and require different quality control measures.**

**(a1)** Representative LXRT images of first and second dorsal slices from an OB sample with ‘patchy’ artefact. **(a2)** Percentage of samples with ‘patchy’ in experiments fixed by immersion or perfusion. Each dot represents an experiment. Error bars show mean +/- standard error of mean. **(b)** Representative SBEM images of granule cells and synapses in the external plexiform layer of OB samples fixed by immersion or perfusion. Arrowheads point towards membrane breaks. **(c)** Structures of SPPs using immersion fixation **(c1)** and perfusion fixation **(c2)**, respectively. Black text in grey boxes states the imaging techniques; white text in boxes states the sample processing steps; diamonds represent quality control steps; grey circles represent curated datasets; blue lines represent correlations among curated datasets; artefacts are shown in coloured boxes next to their hypothetical causes in line with the sample processing step that putatively generates them. Inspection of most SPP artefacts occurs after LXRT. Inspection methods include LXRT (green line) and SBEM (red line).

a

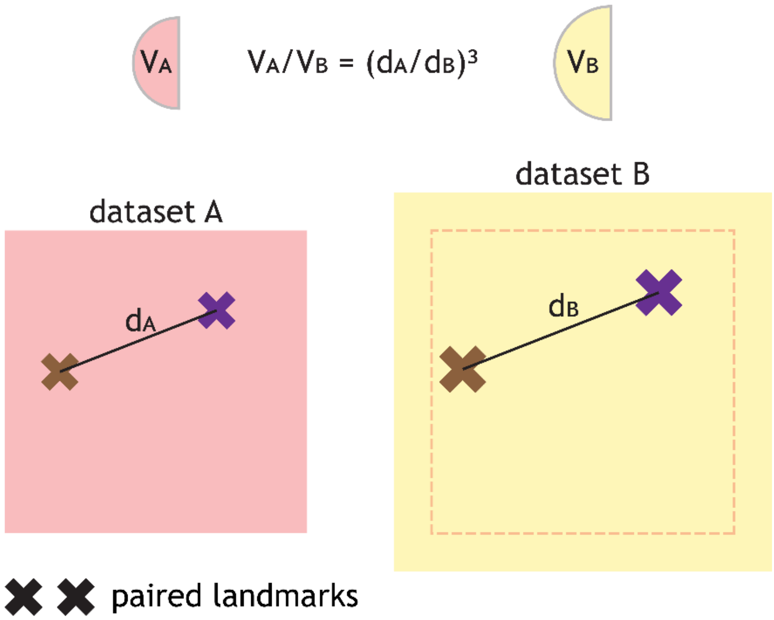


**Supp. F4. Method to estimate the sample volume of fresh tissue samples.**
**(a)** The volume ratio between two volumetric datasets A and B mapping the same specimen is estimated to be the cubed ratio of the distances between conserved pairs of landmarks in both datasets. The dashed box represents the volume of dataset A in the expanded dataset B. Symbols: V: volume, d: distance. This approach was used to estimate the fresh tissue volume of a 1st OB dorsal slice from correlated 2P and SBEM landmarks. The 2P dataset was obtained in the fresh tissue *in vivo*, whereas the SBEM dataset was acquired after staining and embedding had taken place.

**Supp. Table 1. Artefact definition.**
Definitions of each of the 12 artefacts (**Fig. 1c**), subtypes of 'sideways' (**Fig. 2b**) and subtypes of 'crack' (**Fig. 3a1**).

**Supp. Table 2. Sample preparation protocol (SPP) details.**
Summary of samples, and details on the fixation, staining, dehydration and embedding steps of all SPPs included in this study.

**Supp. Table 3. Experimental and animal details of images in figures.**
Animal details, experiment ID and relevant sample details (slice index, thickness, in vivo two-photon history) for every 2P, LXRT, SXRT and EM image shown in the figures.

**Supp. Table 4. Mouse line details.**
Founder line details of all mouse lines used in this study.

**Supp. Table 5. Experimental cohort descriptions.**
Sample IDs and sample selection criteria for all the analyses included in this study.

**Supp. Table 6. Volume cohorts.**
List of samples whose final in-resin volume was measured.
